# Causal relationship of cerebrospinal fluid biomarkers with the risk of Alzheimer’s disease: A two-sample Mendelian randomization study

**DOI:** 10.1101/719898

**Authors:** Soyeon Kim, Kiwon Kim, Kwangsik Nho, Woojae Myung, Hong-Hee Won

**Affiliations:** Samsung Advanced Institute for Health Sciences and Technology (SAIHST), Sungkyunkwan University, Samsung Medical Center, Seoul, South Korea; Department of Psychiatry, Veteran Health Service Medical Center, Seoul, South Korea; Department of Radiology & Imaging Sciences, Indiana University School of Medicine, Indianapolis, IN, USA; Department of Neuropsychiatry, Seoul National University Bundang Hospital, Seongnam, South Korea

**Keywords:** Alzheimer’s disease, CSF, amyloid beta, tau protein, Mendelian randomization

## Abstract

Whether the epidemiological association of amyloid beta (Aβ) and tau pathology with Alzheimer’s disease (AD) is causal remains unclear. The recent failures to demonstrate the efficacy of several amyloid beta-modifying drugs may indicate the possibility that the observed association is not causal. These failures also led to efforts to develop tau-directed treatments whose efficacy is still tentative. Herein, we conducted a two-sample Mendelian randomization analysis to determine whether the relationship between the cerebrospinal fluid (CSF) biomarkers for amyloid and tau pathology and the risk of AD is causal. We used the summary statistics of a genome-wide association study (GWAS) for CSF biomarkers (Aβ_1-42_, phosphorylated tau _181_ [p-tau], and total tau [t-tau]) in 3,146 individuals and for late-onset AD (LOAD) in 21,982 LOAD cases and 41,944 cognitively normal controls. We tested the association between the change in the genetically predicted CSF biomarkers and LOAD risk. We found a modest decrease in the LOAD risk per one standard deviation (SD) increase in the genetically predicted CSF Aβ (odds ratio [OR], 0.63 for AD; 95% confidence interval [CI], 0.38-0.87; *P* = 0.02). In contrast, we observed a significant increase in the LOAD risk per one SD increase in the genetically predicted CSF p-tau (OR, 2.37; 95% CI, 1.46-3.28; *P* = 1.09×10^−5^). However, no causal association was observed of the CSF t-tau with the LOAD risk (OR, 1.15; 95% CI, 0.85-1.45; *P* = 0.29). Our findings need to be validated in future studies with more genetic variants identified in larger GWASs for CSF biomarkers.

## Introduction

Alzheimer’s disease (AD), a leading cause of dementia, is the largest burden source of morbidity and mortality in older adults. One in every 85 individuals is expected to develop AD, which means that delaying the onset by one year can reduce the number of patients with AD worldwide up to 9 million by 2020^1^. Given that eightfold as many individuals have preclinical AD at risk of progression^2^, the development of disease-modifying therapies is urgently required. Amyloid beta (Aβ) peptides are transmembrane amyloid precursor proteins^3^ and tau is a microtubule-associated protein^4^. Decades of research have accumulated the evidence on the pathophysiology of Aβ and tau proteins that independently form plaques and tangles and lead normal functional neurons into a disabled state, AD^5^. Understanding AD as the result of abnormal proteins, extracellular amyloid plaques, and intraneuronal neurofibrillary tau tangles, two-thirds of the novel treatment pipelines aim at disease-modifying therapies, 90% of which are anti-amyloid and anti-tau protein agents^6^.

However, numerous trials to develop novel therapies targeting the amyloid plaques to modify the disease progress recently turned out failures. These failures could bring a reasonable doubt about the role of Aβ in the pathophysiology of AD with delicate elaboration^7^. One possible explanation of the failure of clinical trials targeting the amyloid plaques is that the intervention is performed too late in the disease course to reverse the pathology in the trial participants.^8–10^. However, the poor efficacy of the amyloid-targeting therapy may be due to the amyloid being a downstream result, rather than a cause of AD^11, 12^. With these recent failures, tau protein has gained more attention as a target for disease-modifying therapies. Although previous animal studies showed that the suppression of tau gene expression was protective to cognitive impairment, this impact required accompanying regulation of Aβ^13^. In addition, recent studies of the association between premortem cognitive function and AD neuropathology, including tau protein, have shown vague results^14, 15^. These results also brought on a doubt on the tau pathology in AD^16^. Thus, further research is still required to determine whether Aβ or tau proteins are causal to AD or are surrogate markers for AD. This issue is crucial for the successful development of disease-modifying drugs.

One promising approach for investigating the causality is Mendelian randomization (MR) using genetic variants as the instrumental variables (IVs)^17^. The association between the genetic variants and the disease outcome can provide evidence of causation while, subject to certain assumptions, minimizing confounding factors, including age, education, or other environmental exposures. This method may be useful to elaborate the causal relationship of Aβ or tau protein with AD without confounding factors and reverse causality^18–20^.

Herein, we hypothesized that Aβ or tau protein have a causal effect on the risk for late-onset AD (LOAD), and tested the hypothesis using two-sample MR (TSMR) methods with a summary statistics from large-scale genome-wide association studies (GWASs) of cerebrospinal fluid (CSF) biomarkers (Aβ_1-42_ [Aβ], phosphorylated tau _181_ [p-tau], and total tau [t-tau]) and late-onset AD^21, 22^.

## Materials and methods

### Exposure

In this study, we used three CSF biomarkers for AD, Aβ, p-tau, and t-tau, as exposures for investigating the causal relationship with the outcome of interest. Meta-analyzed GWAS summary statistics of these biomarkers were obtained from 3,146 individuals in nine different studies (Knight ADRC, the Charles F. and Joanne Knight Alzheimer’s Disease Research Center; ADNI1, Alzheimer’s Disease Neuroimaging Initiative phase 1; ADNI2, Alzheimer’s Disease Neuroimaging Initiative phase 2; BIOCARD, Predictors of Cognitive Decline Among Normal Individuals; HB, Saarland University in Homburg/Saar, Germany; MAYO, Mayo Clinic; SWEDEN, Skåne University Hospital; UPENN, Perelman School of Medicine at the University of Pennsylvania; UW, the University of Washington)^21^. The sample size of these GWASs is the largest at present with respect to Aβ, p-tau, and t-tau collected from CSF. The effect per single-nucleotide polymorphism (SNP) in the GWAS summary statistics was defined as a standardized beta coefficient since each phenotype was converted using a log-transformation to follow the normal distribution.

### Outcome

Our outcome of interest was LOAD, defined as AD with an onset at 65 years of age or older. We utilized the summary-level data from the stage 1 meta-analysis of the GWASs for LOAD in the National Institute on Aging Genetics of Alzheimer’s Disease Data Storage Site^22^. The meta-analysis result was obtained from the four consortia (The Alzheimer Disease Genetics Consortium; The European Alzheimer’s disease Initiative; The Cohorts for Heart and Aging Research in Genomic Epidemiology Consortium; and The Genetic and Environmental Risk in AD Consortium Genetic and Environmental Risk in AD/Defining Genetic, Polygenic and Environmental Risk for Alzheimer’s Disease Consortium). It consisted of 46 case-control studies that included 63,926 individuals of European ancestry (21,982 LOAD cases and 41,944 cognitively normal controls).

### Selection of instruments for Mendelian randomization

We performed the following procedures to select appropriate genetic variants that preferentially satisfy three IV assumptions of the MR analysis^23^.

First, we selected the top SNPs with a relaxed threshold (*P* < 1 × 10^−5^), which was considered in recent MR analyses in the case when GWAS for exposure traits only yielded a small number of genome-wide significant SNPs^24–27^. The sample size of the data used in the present study is the largest on CSF biomarkers to this date^21^. CSF biomarkers are expensive, they are acquired through an invasive procedure, and require skilled professionals, which results in a difficulty to gather a sample size sufficient enough to identify many independent SNPs passing a genome-wide significant level (*P* < 5 × 10^−8^)^28^. We relaxed the threshold (*P* < 1 × 10^−5^) to compensate for the small sample size.

Second, we selected the independent genetic variants among those that passed the relaxed threshold, using the cutoff of linkage disequilibrium (LD) value (*r*^2^ < 0.001) to ensure that the IVs for exposure were independent^29^. The LD between the SNPs was calculated based on the European individuals from the 1000 Genomes Project. If a certain SNP was not available in the summary statistics of the outcome, we substituted that SNP with its LD proxy SNP having a high correlation coefficient (*r*^2^ ≥ 0.8) based on the European ancestry using the LDlink (https://ldlink.nci.nih.gov/). If such LD proxy SNP was not found, the SNP was excluded from the IV set.

Third, we eliminated the SNPs that had ambiguous alleles from the IV set when the alleles in the exposure and the outcome were not identical. For example, we excluded an SNP if the effect allele and the non-effect allele of the exposure and outcome were T/C, and T/G, respectively^29^.

Fourth, to ensure that there was no horizontal pleiotropy among the IVs, we conducted an MR-Pleiotropy Residual Sum and Outlier (MR-PRESSO) test that detects pleiotropic variants among the exposure-associated variants^30^. Considering the SNPs that had a direct effect on LOAD, which means a direct pleiotropic effect on the outcome of interest, we excluded rs769449 in the apolipoprotein E (*APOE*) region that is highly associated with LOAD from the set of IVs^31, 32^. The *APOE* region has been reported to have multiple pleiotropic effects in many previous studies^33^. When the MR analysis is performed with the outliers detected by MR-PRESSO or variants in the *APOE* region, including the pleiotropic SNPs in the instruments, it may result in a positive bias or a negative bias due to horizontal pleiotropy and induce inaccurate causal relationship^34^. Therefore, we excluded the outliers detected by MR-PRESSO. Subsequently, to confirm the absence of horizontal pleiotropy, we performed the MR-Egger intercept test with the intercept unconstrained^35^. The intercept of the MR-Egger regression represents a statistical estimate of the directional pleiotropic effect, which can be a confounding factor in MR. The selected genetic variants are listed in **Supplementary Tables 1-3**.

### Two-sample Mendelian randomization method

TSMR utilizes the GWAS summary statistics obtained from two large sample sets, allowing to use more robustly associated genetic instruments compared with one-sample MR^17^. TSMR in the present study was performed using the Two Sample MR R package (version 0.4.22) from the MR-Base platform^29^. To confirm that the findings of the estimation of the causal effect of the exposures on the risk of LOAD are credible, we used diverse methods, including the inverse-variance weighted (IVW), maximum likelihood, weighted median, and MR-Egger regression. These multiple methods have been developed and differ from each other in terms of sensitivity to heterogeneity, bias, and power. We selected the IVW method as our primary MR method because it provides reliable results in the presence of heterogeneity in an MR analysis and is appropriate when using a large number of SNPs^36^. The standard error (SE) of the IVW effect was estimated using a multiplicative random effects model. We performed a leave-one-out analysis that estimates the causal effect of all but one SNP at a time iteratively, using the IVW method, to test if the results were derived from any particular SNP.

The maximum likelihood method is a likelihood-based method that assumes a bivariate normal distribution of the exposure and outcome, which better elucidates the correlation between two different GWAS summary statistics than does the IVW^23^. We also used the weighted median and MR-Egger regression. Since these two methods provide convincing causal estimates in the presence of violation of the MR assumptions, they were used as sensitivity analyses in the MR studies^35, 37^.

We used a forest plot to visualize the heterogeneity between the instruments due to horizontal pleiotropy and the contribution of each instrument to the overall estimate^29^. A funnel plot showing the proportion of the precision (1/ SE) to Wald ratios per SNP was used to evaluate the bias due to the invalid instruments. The overall symmetry in the funnel plot represents the lack of severe heterogeneity and bias driven by directional horizontal pleiotropy that violates the MR assumptions^38^.

### Power calculation

We calculated the statistical power of the MR using an online tool (https://sb452.shinyapps.io/power/)^39^ based on the proportion of variance in the exposure (*R*^2^) explained by genetic instruments, true causal effect of the exposure on the outcome, sample size, and ratio of cases to controls of the outcome. *R*^2^ was obtained from the MR-Steiger directionality test^40^. We estimated the true causal effect based on the observed odds ratios (ORs) between the CSF biomarkers and the risk of LOAD.

## Results

In our main analysis, we excluded the outliers using MR-PRESSO to select suitable instruments that satisfied one of the core IV assumptions, that is, no horizontal pleiotropy. The number of the outlier SNPs predicted to have pleiotropy by MR-PRESSO was one (out of 15 top SNPs) and thirteen (out of 20 top SNPs) for CSF Aβ and p-tau, respectively. After excluding the outliers, the MR-Egger intercept test showed no evidence of horizontal pleiotropy in both Aβ and p-tau (Aβ: intercept = −0.027, SE = 0.015, *P* = 1; p-tau: intercept = −0.004, SE = 0.044, *P* = 0.93) (**Supplementary Table 4**).

CSF Aβ showed evidence for a causal effect on the risk for LOAD (IVW OR, 0.63 for LOAD per 1 standard deviation (SD) increase in the genetically predicted CSF Aβ; 95% confidence interval [CI], 0.38-0.87; *P* = 0.02) (**Table 1 and Fig. 1A**). We found a more prominent causality between CSF p-tau and the risk for LOAD (IVW OR, 2.37 for LOAD per 1 SD increase in the genetically predicted CSF p-tau; 95% CI, 1.46-3.28; *P* = 1.09×10^−5^) (**Table 1 and Fig. 1C**). In the sensitivity analyses, the causal effect of CSF p-tau on the risk for LOAD was significant in both the maximum likelihood and the weighted median (maximum likelihood *P* = 2.98×10^−5^ and weighted median *P* = 2.51×10^−4^), while the causal association between CSF Aβ and the risk for LOAD was significant in the maximum likelihood (*P* = 0.03). Both CSF Aβ and CSF p-tau were not significant in the MR-Egger regression. Although the effect estimate for CSF p-tau was similar to that derived by the IVW, maximum likelihood, and weighted median, the estimates for CSF Aβ were different. MR-Egger was shown to yield minimally biased estimates regardless of the pleiotropic SNPs in the instruments^35^. However, in our MR analysis, the potential pleiotropic SNPs were detected as outliers using the MR-PRESSO and all of them were excluded, which may suggest that the IVW has a greater power and derives a more precise estimate than the MR-Egger regression.

**Table 1.** Two-sample Mendelian randomization for the causal relationship of amyloid beta, phosphorylated tau, and total tau with the risk for late-onset Alzheimer’s disease.

| Exposure | Method | OR (95% CI)<sup>a</sup> | *P* value | No. of SNPs<sup>b</sup> |
| --- | --- | --- | --- | --- |
| Amyloid beta | IVW<sup>c</sup> | 0.63 (0.38 to 0.87) | 0.02 | 14 |
|  | Maximum likelihood<sup>c</sup> | 0.63 (0.38 to 0.89) | 0.03 | 14 |
|  | Weighted median<sup>c</sup> | 0.61 (0.26 to 0.95) | 0.09 | 14 |
|  | MR-Egger<sup>c</sup> | 1.63 (-0.18 to 3.44) | 0.41 | 14 |
| Phosphorylated tau | IVW<sup>c</sup> | 2.37 (1.46 to 3.28) | 1.09×10<sup>-5</sup> | 7 |
|  | Maximum likelihood<sup>c</sup> | 2.39 (1.41 to 3.37) | 2.98×10<sup>-5</sup> | 7 |
|  | Weighted median<sup>c</sup> | 2.50 (1.27 to 3.73) | 2.51×10<sup>-4</sup> | 7 |
|  | MR-Egger<sup>c</sup> | 2.67 (-3.75 to 9.08) | 0.46 | 7 |
| Total tau | IVW<sup>c</sup> | 1.15 (0.85 to 1.45) | 0.29 | 18 |
|  | Maximum likelihood<sup>c</sup> | 1.16 (0.86 to 1.45) | 0.27 | 18 |
|  | Weighted median<sup>c</sup> | 1.14 (0.73 to 1.56) | 0.46 | 18 |
|  | MR-Egger<sup>c</sup> | 2.03 (0.35 to 3.70) | 0.11 | 18 |

Abbreviations: IVW, inverse variance-weighted; MR, Mendelian randomization; OR, odds ratio; CI, confidence interval; SNP, single-nucleotide polymorphism.
<sup>a</sup> Indicates odds for AD per 1 standard deviation increase in amyloid beta, phosphorylated tau, or total tau.
<sup>b</sup> Top SNPs with $P < 1 \times 10^{-5}$ were included in the analysis.
<sup>c</sup> Indicates models without MR-PRESSO (Pleiotropy Residual Sum and Outlier).

**Figure 1.**
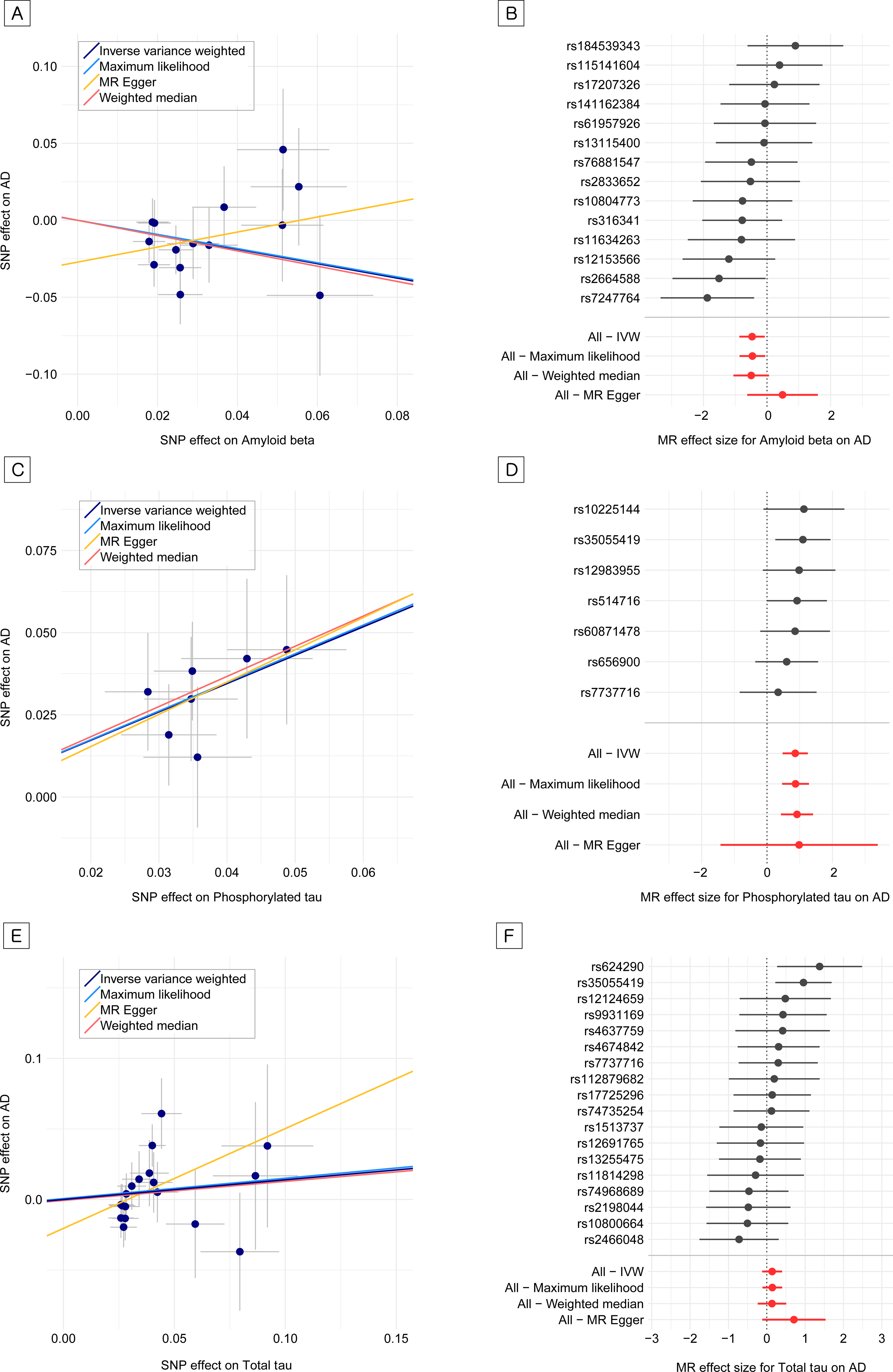
Estimated causal effects of amyloid beta, phosphorylated tau, and total tau on the risk for late-onset Alzheimer’s disease **A.** Scatter plot of the effect size for amyloid beta and the risk for Alzheimer’s disease per genetic variant. **B.** Forest plot of the estimate of amyloid beta on the risk for Alzheimer’s disease for each or all variants. **C.** Scatter plot of the effect size for phosphorylated tau and the risk for Alzheimer’s disease per genetic variant. **D.** Forest plot of the estimate of phosphorylated tau on the risk for Alzheimer’s disease for each or all variants. **E.** Scatter plot of the effect size for total tau and the risk for Alzheimer’s disease per genetic variant. **F.** Forest plot of the estimate of total tau on the risk for Alzheimer’s disease for each or all variants.

While the effect size for CSF Aβ on the risk for LOAD yielded patterns indicating a moderate heterogeneity among the instruments, the instruments of CSF p-tau showed little heterogeneity (**Fig. 1B and Fig. 1D**). The Cochran Q statistics, *P* value, and heterogeneity (*I*^2^ [%]) were 13.41, 0.42, and 3 for CSF Aβ, and 1.58, 0.95, and 0 for CSF p-tau, respectively (**Supplementary Tables 4**). The leave-one-out analysis confirmed that a single SNP was not exclusively responsible for the associations with the risk for LOAD (**Fig. 2A and Fig. 2C**). In the funnel plot, each dot shows the proportion of the precision (1/SE) to Wald ratio per SNP and the vertical lines represent the MR estimates jointed by the instruments (**Fig. 2B and Fig. 2D**). We observed an overall symmetry in the funnel plot, which indicates that our results were less likely biased by invalid instruments.

**Figure 2.**
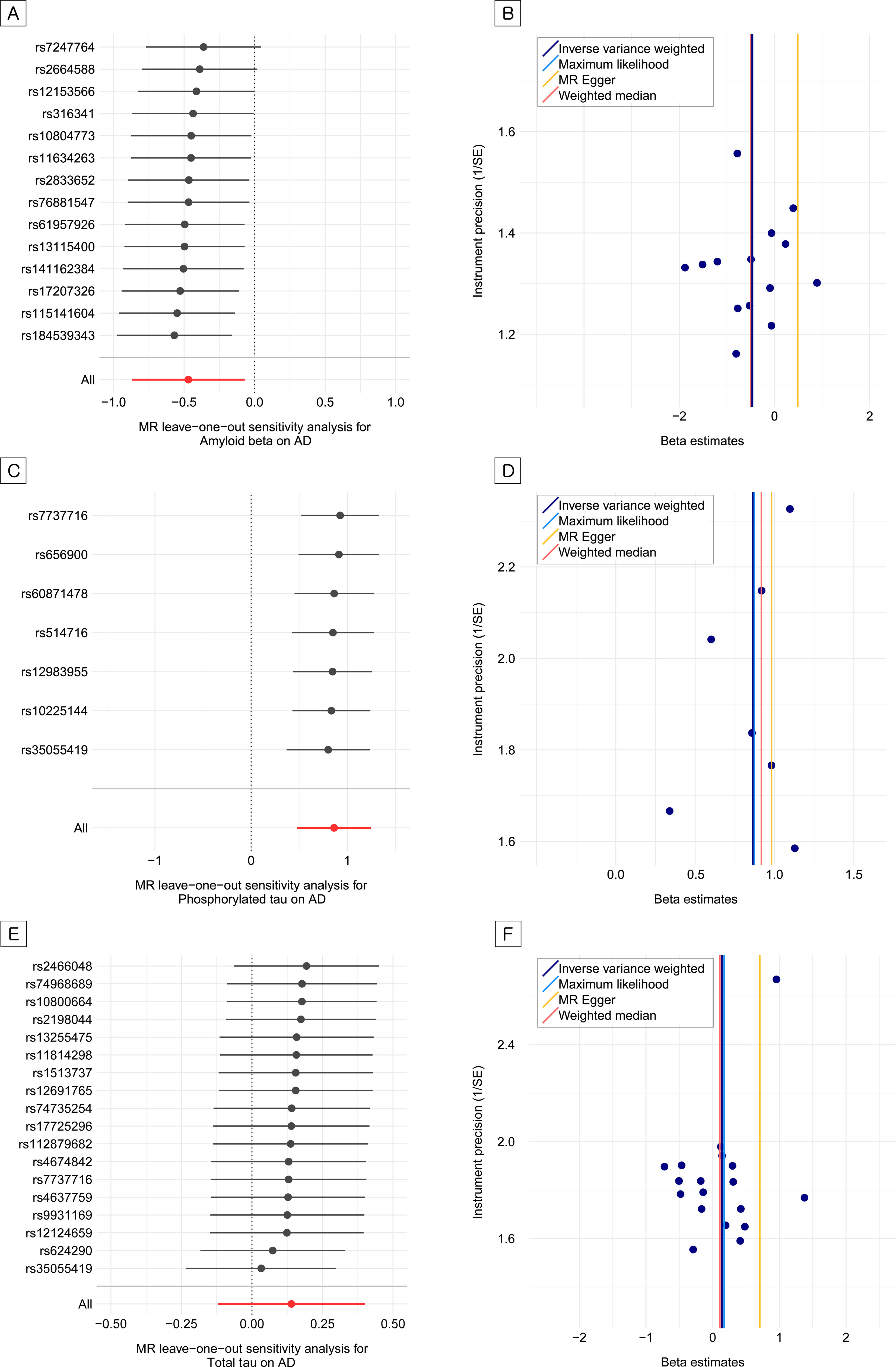
Leave-one-out analyses and funnel plots for amyloid beta, phosphorylated tau, and total tau on the risk for late-onset Alzheimer’s disease **A.** Leave-one-out analysis for amyloid beta on the risk for Alzheimer’s disease. **B.** Funnel plot for amyloid beta on the risk for Alzheimer’s disease. **C.** Leave-one-out analysis for phosphorylated tau on the risk for Alzheimer’s disease. **D.** Funnel plot for phosphorylated tau on the risk for Alzheimer’s disease. **E.** Leave-one-out analysis for total tau on the risk for Alzheimer’s disease. **F.** Funnel plot for total tau on the risk for Alzheimer’s disease.

We also investigated the association between t-tau and the risk for LOAD. In contrast to CSF p-tau, no causal evidence was found for the effect of CSF t-tau on the risk for LOAD (IVW OR, 1.15 for LOAD per 1 SD increase in the genetically predicted CSF t-tau; 95% CI, 0.85-1.45; *P* = 0.29) (**Table 1**, **Fig. 1E**, **Fig. 1F**, **Fig. 2E**, **and Fig. 2F**). There was no outlier detected by MR-PRESSO among the instruments of CSF t-tau. We confirmed that there was no evidence for horizontal pleiotropy (Intercept = −0.020, SE = 0.014, *P* = 0.18) and little heterogeneity between the IVs (Q = 18.45, *P* = 0.36, *I*^2^ [%] = 8) (**Supplementary Table 4**).

Given the observed ORs between the measured CSF biomarkers and the risk for LOAD, our MR analysis showed sufficient statistical power (> 90%) to detect the causal effects of the CSF biomarkers on the risk for LOAD with a level of significance of 0.05. **Supplementary Table 5** presents the estimates of the statistical power for our MR analysis.

## Discussion

Using TSMR with genetic instruments from large-scale GWASs, we investigated the potential causal relationship between CSF biomarkers and the risk for LOAD. In this MR study, the genetic association of the CSF Aβ and p-tau instruments supported the causality of CSF Aβ and p-tau on the risk for LOAD. In contrast, we found no significant evidence of a causal relationship between CSF t-tau and the risk for LOAD. Our results are consistent with those of recent reports ^41–43^.

Although Aβ, p-tau, and t-tau in the CSF have been reported to be useful as disease progression markers^41, 44^, there is still little evidence for their causal relationship with LOAD in randomized clinical trials (RCTs)^45, 46^. Recent RCTs on the elimination of the accumulated Aβ or tau proteins could not provide solid evidence for improvement of the symptoms of LOAD^47–49^. While clinical trials with small sample sizes have shown that eliminating the Aβ elements led to symptomatic improvement^50^, larger studies have failed to establish consistent results^47, 51, 52^. The agents reducing tau phosphorylation represented promising benefits in pilot clinical studies^53, 54^, but failed to show significant improvements in a cohort study^54^; tau aggregation inhibitors showed a similar pattern^55^. Although another approach for proving the causality for LOAD is the induced pathologic accumulation of Aβ and tau proteins in RCTs, such intervention in humans is not allowed due to ethical issues. Instead, the development of AD phenotypes has been attempted in numerous animal models with accumulating Aβ^56, 57^ and tau proteins^58^, and these still have various limitations. Transgenic animal models generally represent familiar AD rather than sporadic LOAD due to targeting a specific pathologic substance; therefore, they cannot provide a full explanation of LOAD^59^. In addition, animal models could not represent the complex symptomatology of dementia that presents in humans.

In consideration of these perspectives, the principles of MR can be applied to provide clues for the causality of these biomarkers in the etiology of LOAD^60^. This approach, which is conceptually similar to that of RCTs,^61^ is based on the Mendel’s law of segregation that genetic variants are randomly allocated at meiosis and that these genetic variants are consequently independent of many confounding factors or reverse causation. Thus, an MR analysis could enable the inference of the risk for LOAD driven through the genetically determined risk of amyloid accumulation and tau pathology. In our study, we found evidence supporting the potential causal relationships between Aβ and p-tau proteins in the CSF and the risk for LOAD using MR with genetic instruments selected from large-scale GWASs.

The causal estimates in our analysis were based on the largest GWAS to date, which may increase the precision of the estimates. We estimated a 0.63-fold decrease of the risk for LOAD per 1 SD increase in the CSF Aβ, and a 2.37-fold increase in the risk for LOAD per 1 SD increase in the CSF p-tau. These directions of association are consistent with those in previous reports^41, 62^. Markedly increased levels of p-tau proteins and decreased levels of Aβ in the CSF are represented as a specific finding in LOAD^41^. Aβ accumulation in the neuronal plaques and its binding to various receptors have been known as a hallmark of LOAD. Aβ binding to receptors has been understood as a process leading to neuronal toxicity, inducing mitochondrial dysfunction and oxidative stress^63, 64^. The pathologic process of tau in LOAD consists of the development of phosphorylated pre-tangles and formation of the neuropil threads^65^. After a process of hyperphosphorylation, acetylation, N-glycosylation, and truncation, tau forms the tangles in LOAD^42^. The causal relationship of Aβ and p-tau observed in our MR analysis supports that Aβ and p-tau may play important roles in the pathophysiology of LOAD. Further studies investigating the biological mechanisms are needed.

T-tau in the CSF and the risk for LOAD did not show a causal relationship, which is consistent with the findings of previous studies^66, 67^. While the CSF level of p-tau increases specifically in LOAD, the CSF level of t-tau can increase in various conditions of neurodegeneration, including LOAD and other brain disorders^68^. Our result may support a recent proposal emphasizing the tau hyperphosphorylation in AD versus the excessive production of tau proteins^42^.

The measured CSF biomarkers in AD reflect both the production and the clearance of these markers at a given time. In contrast, neuroimages represent the neuropathologic load or damage accumulated over time directly in the brain^69^. Thus, imaging GWAS, such as amyloid or tau deposition in the brain measured by positron emission tomography (PET)^70^ as phenotypes, could provide additional information for the association between these biomarkers and the risk for LOAD. However, the sample size of the current imaging genetic studies for these biomarkers is limited. Further studies with larger samples of genetic and imaging data could be helpful.

This study has several limitations. First, our causal estimates may be affected by several factors; horizontal pleiotropy, which was not detected by the applied MR sensitivity analysis methods^71^, and the possibility of misclassified LOAD cases^72, 73^. Unlike the balanced or positive bias induced by horizontal pleiotropy, the misclassified cases in the outcome may lead our results toward null. However, the estimates were statistically significant and consistent in various methods applied in our analysis. Second, our GWAS data included samples of Caucasian ancestry, which may limit the generalization of our findings. Finally, even though we employed the summary statistics from the largest GWASs on Aβ and tau proteins to date^21^, we applied a relaxed threshold to include more IVs as done in other psychiatric MR studies ^24–27^. Despite using instruments with a less stringent threshold, which may lead to null findings, our power analysis of the MR showed a statistical power greater than 90% and our analysis derived significant causal estimates.

In conclusion, this MR analysis suggested a possible causal relationship of the CSF Aβ and p-tau with the risk for LOAD. In addition, our findings showed that the association between t-tau and the risk for LOAD was not causal. Our results suggested that the etiology of LOAD involves multiple biological processes, including the amyloid and tau proteins in the AD pathophysiology. This complex nature of LOAD could partly explain the recent multiple failures of clinical trials of anti-amyloid monotherapy^47, 51, 52, 74, 75^. Further MR studies for multiple candidate biomarkers could be helpful to find appropriate drug targets for LOAD and larger GWAS data with sufficient numbers of IVs are necessary to validate the causality of CSF Aβ and p-tau on the risk for LOAD.

## Acknowledgments

The authors thank the researchers at the Washington University School of Medicine for providing the summary statistics of GWAS for CSF biomarkers.

This work was supported by the a grant from the National Research Foundation (NRF) funded by the Ministry of Science and ICT (MSIT) [grant number NRF-2019R1A2C4070496 to HH Won; NRF-2018R1C1B6001708 to W Myung]. This work was also supported by grants from the National Institutes of Health [grant number R01LM012535, R03AG054936, and R03AG063250 to K Nho].

## References

1. Brookmeyer R, Johnson E, Ziegler-Graham K, Arrighi HM. Forecasting the global burden of Alzheimer’s disease. Alzheimers Dement 2007; 3(3): 186–191.

2. Brookmeyer R, Abdalla N, Kawas CH, Corrada MM. Forecasting the prevalence of preclinical and clinical Alzheimer’s disease in the United States. Alzheimers Dement 2018; 14(2): 121–129.

3. Chen GF, Xu TH, Yan Y, Zhou YR, Jiang Y, Melcher K et al. Amyloid beta: structure, biology and structure-based therapeutic development. Acta Pharmacol Sin 2017; 38(9): 1205–1235.

4. Pradeepkiran JA, Reddy PH. Structure Based Design and Molecular Docking Studies for Phosphorylated Tau Inhibitors in Alzheimer’s Disease. Cells 2019; 8(3).

5. Bloom GS. Amyloid-beta and tau: the trigger and bullet in Alzheimer disease pathogenesis. JAMA Neurol 2014; 71(4): 505–508.

6. Cummings J, Lee G, Ritter A, Zhong K. Alzheimer’s disease drug development pipeline: 2018. Alzheimers Dement (N Y) 2018; 4: 195–214.

7. van Dyck CH. Anti-Amyloid-beta Monoclonal Antibodies for Alzheimer’s Disease: Pitfalls and Promise. Biol Psychiatry 2018; 83(4): 311–319.

8. Villemagne VL, Pike KE, Chetelat G, Ellis KA, Mulligan RS, Bourgeat P et al. Longitudinal assessment of Abeta and cognition in aging and Alzheimer disease. Ann Neurol 2011; 69(1): 181–192.

9. Jack CRJr., Wiste HJ, Lesnick TG, Weigand SD, Knopman DS, Vemuri P et al. Brain beta-amyloid load approaches a plateau. Neurology 2013; 80(10): 890–896.

10. Knopman DS, Jack CRJr., Lundt ES, Weigand SD, Vemuri P, Lowe VJ et al. Evolution of neurodegeneration-imaging biomarkers from clinically normal to dementia in the Alzheimer disease spectrum. Neurobiol Aging 2016; 46: 32–42.

11. Sagare AP, Bell RD, Zlokovic BV. Neurovascular defects and faulty amyloid-beta vascular clearance in Alzheimer’s disease. J Alzheimers Dis 2013; 33 Suppl 1: S87–100.

12. Zlokovic BV. Neurovascular mechanisms of Alzheimer’s neurodegeneration. Trends Neurosci 2005; 28(4): 202–208.

13. Busche MA, Wegmann S, Dujardin S, Commins C, Schiantarelli J, Klickstein N et al. Tau impairs neural circuits, dominating amyloid-beta effects, in Alzheimer models in vivo. Nat Neurosci 2019; 22(1): 57–64.

14. Silver MH, Newell K, Brady C, Hedley-White ET, Perls TT. Distinguishing between neurodegenerative disease and disease-free aging: correlating neuropsychological evaluations and neuropathological studies in centenarians. Psychosom Med 2002; 64(3): 493–501.

15. Berlau DJ, Corrada MM, Head E, Kawas CH. APOE epsilon2 is associated with intact cognition but increased Alzheimer pathology in the oldest old. Neurology 2009; 72(9): 829–834.

16. Sierksma A, Lu A, Salta E, Vanden Eynden E, Callaerts-Vegh Z, D’Hooge R et al. Deregulation of neuronal miRNAs induced by amyloid-beta or TAU pathology. Mol Neurodegener 2018; 13(1): 54.

17. Davey Smith G, Hemani G. Mendelian randomization: genetic anchors for causal inference in epidemiological studies. Human molecular genetics 2014; 23(R1): R89–98.

18. Lleo A, Alcolea D, Martinez-Lage P, Scheltens P, Parnetti L, Poirier J et al. Longitudinal cerebrospinal fluid biomarker trajectories along the Alzheimer’s disease continuum in the BIOMARKAPD study. Alzheimers Dement 2019.

19. Curtis C, Gamez JE, Singh U, Sadowsky CH, Villena T, Sabbagh MN et al. Phase 3 trial of flutemetamol labeled with radioactive fluorine 18 imaging and neuritic plaque density. JAMA Neurol 2015; 72(3): 287–294.

20. Schonhaut DR, McMillan CT, Spina S, Dickerson BC, Siderowf A, Devous MDSr., et al. (18) F-flortaucipir tau positron emission tomography distinguishes established progressive supranuclear palsy from controls and Parkinson disease: A multicenter study. Ann Neurol 2017; 82(4): 622–634.

21. Deming Y, Li Z, Kapoor M, Harari O, Del-Aguila JL, Black K et al. Genome-wide association study identifies four novel loci associated with Alzheimer’s endophenotypes and disease modifiers. Acta neuropathologica 2017; 133(5): 839–856.

22. Kunkle BW, Grenier-Boley B, Sims R, Bis JC, Damotte V, Naj AC et al. Genetic meta-analysis of diagnosed Alzheimer’s disease identifies new risk loci and implicates Abeta, tau, immunity and lipid processing. Nature genetics 2019; 51(3): 414–430.

23. Burgess S, Small DS, Thompson SG. A review of instrumental variable estimators for Mendelian randomization. Statistical methods in medical research 2017; 26(5): 2333–2355.

24. Gage SH, Jones HJ, Burgess S, Bowden J, Davey Smith G, Zammit S et al. Assessing causality in associations between cannabis use and schizophrenia risk: a two-sample Mendelian randomization study. Psychological medicine 2017; 47(5): 971–980.

25. Choi KW, Chen CY, Stein MB, Klimentidis YC, Wang MJ, Koenen KC et al. Assessment of Bidirectional Relationships Between Physical Activity and Depression Among Adults: A 2-Sample Mendelian Randomization Study. JAMA psychiatry 2019.

26. Hartwig FP, Borges MC, Horta BL, Bowden J, Davey Smith G. Inflammatory Biomarkers and Risk of Schizophrenia: A 2-Sample Mendelian Randomization Study. JAMA psychiatry 2017; 74(12): 1226–1233.

27. Sanna S, van Zuydam NR, Mahajan A, Kurilshikov A, Vich Vila A, Vosa U et al. Causal relationships among the gut microbiome, short-chain fatty acids and metabolic diseases. Nature genetics 2019; 51(4): 600–605.

28. Hansson O, Seibyl J, Stomrud E, Zetterberg H, Trojanowski JQ, Bittner T et al. CSF biomarkers of Alzheimer’s disease concord with amyloid-beta PET and predict clinical progression: A study of fully automated immunoassays in BioFINDER and ADNI cohorts. Alzheimer’s & dementia : the journal of the Alzheimer’s Association 2018; 14(11): 1470–1481.

29. Hemani G, Zheng J, Elsworth B, Wade KH, Haberland V, Baird D et al. The MR-Base platform supports systematic causal inference across the human phenome. eLife 2018; 7.

30. Verbanck M, Chen CY, Neale B, Do R. Detection of widespread horizontal pleiotropy in causal relationships inferred from Mendelian randomization between complex traits and diseases. Nature genetics 2018; 50(5): 693–698.

31. Solovieff N, Cotsapas C, Lee PH, Purcell SM, Smoller JW. Pleiotropy in complex traits: challenges and strategies. Nature reviews Genetics 2013; 14(7): 483–495.

32. Lambert JC, Ibrahim-Verbaas CA, Harold D, Naj AC, Sims R, Bellenguez C et al. Meta-analysis of 74,046 individuals identifies 11 new susceptibility loci for Alzheimer’s disease. Nature genetics 2013; 45(12): 1452–1458.

33. Tuminello ER, Han SD. The apolipoprotein e antagonistic pleiotropy hypothesis: review and recommendations. International journal of Alzheimer’s disease 2011; 2011: 726197.

34. Holmes MV, Ala-Korpela M, Smith GD. Mendelian randomization in cardiometabolic disease: challenges in evaluating causality. Nature reviews Cardiology 2017; 14(10): 577–590.

35. Bowden J, Davey Smith G, Burgess S. Mendelian randomization with invalid instruments: effect estimation and bias detection through Egger regression. International journal of epidemiology 2015; 44(2): 512–525.

36. Burgess S, Butterworth A, Thompson SG. Mendelian randomization analysis with multiple genetic variants using summarized data. Genetic epidemiology 2013; 37(7): 658–665.

37. Bowden J, Davey Smith G, Haycock PC, Burgess S. Consistent Estimation in Mendelian Randomization with Some Invalid Instruments Using a Weighted Median Estimator. Genet Epidemiol 2016; 40(4): 304–314.

38. Sterne JA, Sutton AJ, Ioannidis JP, Terrin N, Jones DR, Lau J et al. Recommendations for examining and interpreting funnel plot asymmetry in meta-analyses of randomised controlled trials. BMJ (Clinical research ed) 2011; 343: d4002.

39. Burgess S. Sample size and power calculations in Mendelian randomization with a single instrumental variable and a binary outcome. International journal of epidemiology 2014; 43(3): 922–929.

40. Hemani G, Tilling K, Davey Smith G. Orienting the causal relationship between imprecisely measured traits using GWAS summary data. PLoS genetics 2017; 13(11): e1007081.

41. Olsson B, Lautner R, Andreasson U, Ohrfelt A, Portelius E, Bjerke M et al. CSF and blood biomarkers for the diagnosis of Alzheimer’s disease: a systematic review and meta-analysis. Lancet Neurol 2016; 15(7): 673–684.

42. Congdon EE, Sigurdsson EM. Tau-targeting therapies for Alzheimer disease. Nat Rev Neurol 2018; 14(7): 399–415.

43. Kikuchi K, Kidana K, Tatebe T, Tomita T. Dysregulated Metabolism of the Amyloid-beta Protein and Therapeutic Approaches in Alzheimer Disease. J Cell Biochem 2017; 118(12): 4183–4190.

44. Jack CR, Jr., Wiste HJ, Therneau TM, Weigand SD, Knopman DS, Mielke MM et al. Associations of Amyloid, Tau, and Neurodegeneration Biomarker Profiles With Rates of Memory Decline Among Individuals Without Dementia. Jama 2019; 321(23): 2316–2325.

45. Maass A, Lockhart SN, Harrison TM, Bell RK, Mellinger T, Swinnerton K et al. Entorhinal Tau Pathology, Episodic Memory Decline, and Neurodegeneration in Aging. J Neurosci 2018; 38(3): 530–543.

46. Duyckaerts C, Braak H, Brion JP, Buee L, Del Tredici K, Goedert M et al. PART is part of Alzheimer disease. Acta Neuropathol 2015; 129(5): 749–756.

47. Egan MF, Kost J, Tariot PN, Aisen PS, Cummings JL, Vellas B et al. Randomized Trial of Verubecestat for Mild-to-Moderate Alzheimer’s Disease. N Engl J Med 2018; 378(18): 1691–1703.

48. Egan MF, Kost J, Voss T, Mukai Y, Aisen PS, Cummings JL et al. Randomized Trial of Verubecestat for Prodromal Alzheimer’s Disease. N Engl J Med 2019; 380(15): 1408–1420.

49. Lawlor B, Segurado R, Kennelly S, Olde Rikkert MGM, Howard R, Pasquier F et al. Nilvadipine in mild to moderate Alzheimer disease: A randomised controlled trial. PLoS Med 2018; 15(9): e1002660.

50. Koelsch G. BACE1 Function and Inhibition: Implications of Intervention in the Amyloid Pathway of Alzheimer’s Disease Pathology. Molecules 2017; 22(10).

51. Honig LS, Vellas B, Woodward M, Boada M, Bullock R, Borrie M et al. Trial of Solanezumab for Mild Dementia Due to Alzheimer’s Disease. N Engl J Med 2018; 378(4): 321–330.

52. Relkin NR, Thomas RG, Rissman RA, Brewer JB, Rafii MS, van Dyck CH et al. A phase 3 trial of IV immunoglobulin for Alzheimer disease. Neurology 2017; 88(18): 1768–1775.

53. del Ser T, Steinwachs KC, Gertz HJ, Andres MV, Gomez-Carrillo B, Medina M et al. Treatment of Alzheimer’s disease with the GSK-3 inhibitor tideglusib: a pilot study. J Alzheimers Dis 2013; 33(1): 205–215.

54. Lovestone S, Boada M, Dubois B, Hull M, Rinne JO, Huppertz HJ et al. A phase II trial of tideglusib in Alzheimer’s disease. J Alzheimers Dis 2015; 45(1): 75–88.

55. Gauthier S, Feldman HH, Schneider LS, Wilcock GK, Frisoni GB, Hardlund JH et al. Efficacy and safety of tau-aggregation inhibitor therapy in patients with mild or moderate Alzheimer’s disease: a randomised, controlled, double-blind, parallel-arm, phase 3 trial. Lancet 2016; 388(10062): 2873–2884.

56. Games D, Adams D, Alessandrini R, Barbour R, Berthelette P, Blackwell C et al. Alzheimer-type neuropathology in transgenic mice overexpressing V717F beta-amyloid precursor protein. Nature 1995; 373(6514): 523–527.

57. Hsiao K, Chapman P, Nilsen S, Eckman C, Harigaya Y, Younkin S et al. Correlative memory deficits, Abeta elevation, and amyloid plaques in transgenic mice. Science 1996; 274(5284): 99–102.

58. Andorfer C, Kress Y, Espinoza M, de Silva R, Tucker KL, Barde YA et al. Hyperphosphorylation and aggregation of tau in mice expressing normal human tau isoforms. J Neurochem 2003; 86(3): 582–590.

59. Shinohara M, Fujioka S, Murray ME, Wojtas A, Baker M, Rovelet-Lecrux A et al. Regional distribution of synaptic markers and APP correlate with distinct clinicopathological features in sporadic and familial Alzheimer’s disease. Brain 2014; 137(Pt 5): 1533–1549.

60. Lawlor DA, Harbord RM, Sterne JA, Timpson N, Davey Smith G. Mendelian randomization: using genes as instruments for making causal inferences in epidemiology. Statistics in medicine 2008; 27(8): 1133–1163.

61. Thanassoulis G, O’Donnell CJ. Mendelian randomization: nature’s randomized trial in the post–genome era. Jama 2009; 301(22): 2386–2388.

62. Blennow K, Hampel H. CSF markers for incipient Alzheimer’s disease. Lancet Neurol 2003; 2(10): 605–613.

63. Canevari L, Abramov AY, Duchen MR. Toxicity of amyloid beta peptide: tales of calcium, mitochondria, and oxidative stress. Neurochem Res 2004; 29(3): 637–650.

64. Kim T, Vidal GS, Djurisic M, William CM, Birnbaum ME, Garcia KC et al. Human LilrB2 is a beta-amyloid receptor and its murine homolog PirB regulates synaptic plasticity in an Alzheimer’s model. Science 2013; 341(6152): 1399–1404.

65. Wharton SB, Minett T, Drew D, Forster G, Matthews F, Brayne C et al. Epidemiological pathology of Tau in the ageing brain: application of staging for neuropil threads (BrainNet Europe protocol) to the MRC cognitive function and ageing brain study. Acta Neuropathol Commun 2016; 4: 11.

66. Mattsson N, Zetterberg H, Hansson O, Andreasen N, Parnetti L, Jonsson M et al. CSF biomarkers and incipient Alzheimer disease in patients with mild cognitive impairment. JAMA 2009; 302(4): 385–393.

67. Buerger K, Ewers M, Pirttila T, Zinkowski R, Alafuzoff I, Teipel SJ et al. CSF phosphorylated tau protein correlates with neocortical neurofibrillary pathology in Alzheimer’s disease. Brain 2006; 129(Pt 11): 3035–3041.

68. Ost M, Nylen K, Csajbok L, Ohrfelt AO, Tullberg M, Wikkelso C et al. Initial CSF total tau correlates with 1-year outcome in patients with traumatic brain injury. Neurology 2006; 67(9): 1600–1604.

69. Gordon BA, Friedrichsen K, Brier M, Blazey T, Su Y, Christensen J et al. The relationship between cerebrospinal fluid markers of Alzheimer pathology and positron emission tomography tau imaging. Brain 2016; 139(Pt 8): 2249–2260.

70. Yan Q, Nho K, Del-Aguila JL, Wang X, Risacher SL, Fan KH et al. Genome-wide association study of brain amyloid deposition as measured by Pittsburgh Compound-B (PiB)-PET imaging. Molecular psychiatry 2018.

71. Zheng J, Baird D, Borges MC, Bowden J, Hemani G, Haycock P et al. Recent Developments in Mendelian Randomization Studies. Curr Epidemiol Rep 2017; 4(4): 330–345.

72. Sabbagh MN, Lue LF, Fayard D, Shi J. Increasing Precision of Clinical Diagnosis of Alzheimer’s Disease Using a Combined Algorithm Incorporating Clinical and Novel Biomarker Data. Neurol Ther 2017; 6(Suppl 1): 83–95.

73. Boustani M, Peterson B, Hanson L, Harris R, Lohr KN, Force USPST. Screening for dementia in primary care: a summary of the evidence for the U.S. Preventive Services Task Force. Ann Intern Med 2003; 138(11): 927–937.

74. Ibrahim MM, Gabr MT. Multitarget therapeutic strategies for Alzheimer’s disease. Neural Regen Res 2019; 14(3): 437–440.

75. Morris JC. Editorial: Is Now the Time for Combination Therapies for Alzheimer Disease? The journal of prevention of Alzheimer’s disease 2019; 6(3): 153–154.

