## Supplement for "Causal relationship of cerebrospinal fluid biomarkers with the risk of Alzheimer’s disease: A two-sample Mendelian randomization study"

**SUPPLEMENTARY MATERIALS**

Supplementary Table 1. Genetic instruments of CSF amyloid beta included in the Mendelian randomization analysis

| SNP | Effect allele | Other  allele | Beta for  exposure | Standard error for  exposure | P value for  exposure | Beta for  outcome | Standard error for  outcome | P value for  outcome |
| --- | --- | --- | --- | --- | --- | --- | --- | --- |
| rs316341 | G | A | -0.0246 | 0.004352 | 1.72E-08 | 0.0192 | 0.0158 | 0.2227 |
| rs141162384 | T | G | 0.05123 | 0.010244 | 6.01E-07 | -0.0032 | 0.0366 | 0.9311 |
| rs12153566 | T | A | -0.02566 | 0.005299 | 1.35E-06 | 0.0308 | 0.0191 | 0.1065 |
| rs2664588 | T | C | 0.01913 | 0.004026 | 2.11E-06 | -0.0289 | 0.0143 | 0.04398 |
| rs13115400 | A | G | 0.01924 | 0.004123 | 3.18E-06 | -0.0018 | 0.0149 | 0.9016 |
| rs61957926 | T | C | 0.01874 | 0.004052 | 3.91E-06 | -0.0012 | 0.0154 | 0.9391 |
| rs7247764 | C | T | -0.0257 | 0.005569 | 4.12E-06 | 0.0483 | 0.0193 | 0.01254 |
| rs115141604 | G | A | 0.05535 | 0.012033 | 4.40E-06 | 0.0218 | 0.0382 | 0.5683 |
| rs76881547 | T | C | -0.03289 | 0.007169 | 4.65E-06 | 0.0162 | 0.0244 | 0.5076 |
| rs17207326 | A | G | 0.03666 | 0.008001 | 4.83E-06 | 0.0085 | 0.0266 | 0.7488 |
| rs11634263 | C | T | -0.06063 | 0.013319 | 5.53E-06 | 0.0488 | 0.0522 | 0.3503 |
| rs184539343 | T | C | 0.05141 | 0.011561 | 9.03E-06 | 0.0459 | 0.0395 | 0.2452 |
| rs10804773 | G | A | 0.01789 | 0.004038 | 9.77E-06 | -0.0138 | 0.0143 | 0.3328 |
| rs2833652 | C | T | 0.0289 | 0.006528 | 9.95E-06 | -0.0151 | 0.023 | 0.5122 |

Abbreviations: CSF, cerebrospinal fluid; SNP, single nucleotide polymorphism.

Supplementary Table 2. Genetic instruments of phosphorylated tau included in the Mendelian randomization analysis

| SNP | Effect allele | Other  allele | Beta for  exposure | Standard error for  exposure | P value for  exposure | Beta for  outcome | Standard error for  outcome | P value for  outcome |
| --- | --- | --- | --- | --- | --- | --- | --- | --- |
| rs35055419 | C | T | 0.0349 | 0.005653 | 7.62E-10 | 0.0383 | 0.015 | 0.01083 |
| rs514716 | C | T | -0.04876 | 0.008763 | 2.94E-08 | -0.0448 | 0.0227 | 0.04831 |
| rs60871478 | A | G | 0.03472 | 0.006864 | 4.51E-07 | 0.0298 | 0.0189 | 0.1164 |
| rs10225144 | C | T | 0.02837 | 0.00631 | 7.20E-06 | 0.032 | 0.0179 | 0.07457 |
| rs7737716 | T | C | 0.03566 | 0.007951 | 7.59E-06 | 0.0121 | 0.0214 | 0.5706 |
| rs656900 | C | T | -0.03144 | 0.007018 | 7.97E-06 | -0.0189 | 0.0154 | 0.218 |
| rs12983955 | A | G | -0.04292 | 0.009651 | 9.24E-06 | -0.0421 | 0.0243 | 0.08323 |

Abbreviations: SNP, single nucleotide polymorphism.

Supplementary Table 3. Genetic instruments of total tau included in the Mendelian randomization analysis

| SNP | Effect allele | Other  allele | Beta for  exposure | Standard error for  exposure | P value for  exposure | Beta for  outcome | Standard error for  outcome | P value for  outcome |
| --- | --- | --- | --- | --- | --- | --- | --- | --- |
| rs35055419 | C | T | 0.04004 | 0.006005 | 3.07E-11 | 0.0383 | 0.015 | 0.01083 |
| rs624290 | C | T | -0.04421 | 0.009093 | 1.22E-06 | -0.0609 | 0.025 | 0.01497 |
| rs7737716 | T | C | 0.04066 | 0.008535 | 1.98E-06 | 0.0121 | 0.0214 | 0.5706 |
| rs4674842 | G | T | -0.03081 | 0.00649 | 2.16E-06 | -0.0095 | 0.0168 | 0.5703 |
| rs1513737 | T | C | 0.02597 | 0.005621 | 3.99E-06 | -0.0037 | 0.0145 | 0.8001 |
| rs13255475 | T | C | -0.02793 | 0.006048 | 4.03E-06 | 0.005 | 0.0152 | 0.7431 |
| rs10800664 | T | C | -0.02591 | 0.005654 | 4.77E-06 | 0.0131 | 0.0141 | 0.3549 |
| rs2198044 | A | G | 0.02782 | 0.00609 | 5.11E-06 | -0.0134 | 0.0156 | 0.3897 |
| rs17725296 | C | G | 0.02835 | 0.006205 | 5.13E-06 | 0.004 | 0.0146 | 0.7867 |
| rs12691765 | A | T | -0.02652 | 0.00585 | 6.03E-06 | 0.0044 | 0.0154 | 0.7766 |
| rs2466048 | G | T | 0.02712 | 0.005999 | 6.42E-06 | -0.0196 | 0.0143 | 0.1724 |
| rs112879682 | A | G | -0.08653 | 0.019173 | 6.64E-06 | -0.0168 | 0.0523 | 0.7481 |
| rs11814298 | A | G | -0.05939 | 0.013177 | 6.84E-06 | 0.0174 | 0.0382 | 0.6482 |
| rs74968689 | G | C | 0.07951 | 0.017716 | 7.45E-06 | -0.037 | 0.0418 | 0.3761 |
| rs74735254 | A | C | -0.04236 | 0.009479 | 8.17E-06 | -0.0052 | 0.0214 | 0.8071 |
| rs9931169 | G | A | -0.0341 | 0.007673 | 9.15E-06 | -0.0144 | 0.0198 | 0.4657 |
| rs4637759 | A | G | -0.09193 | 0.020686 | 9.18E-06 | -0.038 | 0.0578 | 0.5106 |
| rs12124659 | C | T | -0.03875 | 0.008755 | 9.95E-06 | -0.0187 | 0.0235 | 0.4265 |

Abbreviations: SNP, single nucleotide polymorphism.

Supplementary Table 4. MR-Egger intercept test for directional horizontal pleiotropy and heterogeneity test.

| Exposure | No. of SNPs | MR-Egger intercept test | | | Heterogeneity test | | |
| --- | --- | --- | --- | --- | --- | --- | --- |
|  |  | Intercept | Standard error | P value | Q (df) | P value | *I*^2^ (%) |
| CSF amyloid beta | 14 | -0.027 | 0.015 | 1 | 13.41 (13) | 0.42 | 3 |
| CSF phosphorylated tau | 7 | -0.004 | 0.044 | 0.93 | 1.58 (6) | 0.95 | 0 |
| CSF total tau | 18 | -0.020 | 0.014 | 0.18 | 18.45 (17) | 0.36 | 8 |

Abbreviations: CSF, cerebrospinal fluid; SNPs, single nucleotide polymorphisms; Q, Cochran's heterogeneity statistic; df, the degrees of freedom; *I*^2^, 100%×(Q - df)/Q^1^.

Supplementary Table 5. Power calculation for Mendelian randomization

|  | Measurement | Exposure > outcome | Outcome  sample size | Ratio of cases to control | Estimated *R*^2 a^ (Number of SNPs) | Causal effect ^b^ | Power |
| --- | --- | --- | --- | --- | --- | --- | --- |
| Without outliers | CSF | Aβ > AD | 63,926 | 1 to 1.91^c^ | 0.101 (14 SNPs) | 0.24 | >90% |
|  | CSF | p-tau > AD | 63,926 | 1 to 1.91^c^ | 0.064 (7 SNPs) | 4.03 | >90% |
| No outliers | CSF | t-tau > AD | 63,926 | 1 to 1.91^c^ | 0.131 (18 SNPs) | 3.67 | >90% |

Abbreviations: SNP, single-nucleotide polymorphism; CSF, cerebrospinal fluid; Aβ, amyloid beta; AD, Alzheimer’s disease; p-tau, phosphorylated tau; t-tau, total tau.

^a^ Coefficient of determination (*R*^2^) of genetic variants on exposure.

^b^ True causal effect (odds ratio, exp(β1)) per 1 standard deviation change in exposure. The estimates were drawn from the population sample set.

^c^ 21,982 Alzheimer’s disease cases and 41,944 cognitively normal controls (in stage 1 Kunkle et al, 2019^2^).

**References**

1.	Higgins JP, Thompson SG, Deeks JJ, Altman DG. Measuring inconsistency in meta-analyses. *BMJ (Clinical research ed)* 2003; **327**(7414)**:** 557-560.

2.	Kunkle BW, Grenier-Boley B, Sims R, Bis JC, Damotte V, Naj AC *et al.* Genetic meta-analysis of diagnosed Alzheimer's disease identifies new risk loci and implicates Abeta, tau, immunity and lipid processing. *Nature genetics* 2019; **51**(3)**:** 414-430.
